# Biogenic phosphonate utilization by globally distributed diatom *Thalassiosira*

**DOI:** 10.1101/2023.08.02.551492

**Authors:** Huilin Shu, Yuan Shen, Hongwei Wang, Xueqiong Sun, Jian Ma, Xin Lin

## Abstract

Phosphonate is a class of enigmatic organic phosphorus compounds that contribute ∼25% of total dissolved organic phosphorus (DOP). Recent studies reveal the important role of phosphonate mediated by prokaryotes in the marine P redox cycle, however, its bioavailability and metabolic process by eukaryotic phytoplankton are under debate. 2-Aminoethylphosphonic acid and (2-AEP) and 2-amino-3-phosphonopropionic acid (2-AP3) are two biogenic phosphonates that ubiquitously exist in marine environment. Here, we report that *Thalassiosira pseudonana*, a dominant diatom species in the ocean, are able to recover growth from phosphorus starvation with individual supplement of 2-AEP and 2-AP3. Moreover, the cellular stoichiometric C:P and N:P ratios of cells grown under 2-AEP are in-between the P-depleted and DIP-replete groups. This study provided evidence that biogenic phosphonates can be adopted as alternative phosphorus sources to support diatom growth and might have a profound effect on the elemental stoichiometry in the oligotrophic ocean.

**Scientific significance statement:** Phosphorus (P) is a major limiting macronutrient for primary productivity in the ocean, whilst the biologically mediated cycling of different P forms is less understood compared to carbon and nitrogen. Accounting for 25% of the marine dissolved organic phosphorus pool, phosphonate bioavailability by eukaryotic phytoplankton still remains discrepancies. In line with our hypothesis, here we report that a cosmopolitan diatom *Thalassiosira pseudonana* can utilize biogenic phosphonates 2-aminoethylphosphonic acid and 2-amino-3-phosphonopropionic acid to support cell growth, with deviated cellular C:N:P in comparison to that of P-replete cells. Furthermore, cells grown under lower temperature exhibit physiological adaptation (*K*-selection strategy) with the benefit of 2-AEP supplied in the medium. We provide the evidence that utilization of biogenic phosphonate is ubiquitous in diatom and it might have profound effect on elemental stoichiometry ratios in the environment. Filling in this missing puzzle piece provides us with a fresh perspective to elucidate the role of phosphonates in marine phosphorus cycle and biological pump.

## Introduction

Phosphorus is an essential nutrient for living organisms, being a vital constituent of phospholipids, nucleic acid, and adenosine triphosphate (ATP). Phytoplankton preferably utilize DIP (dissolved inorganic phosphorus) in the surface ocean. However, low DIP concentration is observed in many regions and has become the major limiting factor for phytoplankton growth and primary productivity [1-3]. In these DIP deficiency regions, DOP (dissolved organic phosphorus) has emerged as a prominent alternative phosphorus source [4, 5].

Phosphonate is a class of organic phosphorus with a characteristic chemically stable C-P bond. Phosphonates in the aquatic environment can be categorized into two types according to the source, chemically synthetic (e.g. herbicide glyphosate) and biogenic. 2-Aminoethylphosphonic acid (2-AEP) and its derivative, 2-amino-3-phosphonopropionic acid (2-AP3) are two biogenic phosphonates which ubiquitously exist in the aquatic ecosystem [6-8]. They are the composition of membrane phospholipids in many organisms, such as bacteria, cyanobacteria and mollusk [9-11]. A recent study shows that *Prochlorococcus* is a major producer of biogenic phosphonate 2-AEP in the ocean, likely allocating over 40% of cellular P towards phosphonate production [10]. Therefore, the bioavailability of biogenic phosphonate represents a prominent alternative phosphorus supply for plankton growth in the open ocean. In comparison with the well-elucidated metabolism pathway in prokaryotic microorganisms [12-14], few attempts have been conducted to explore phosphonate bioavailability to eukaryotic phytoplankton [15, 16]. It has been reported that the picoprasinophyte *Micromonas commode* and coccolithophore *Emiliania huxleyi* are able to utilize 2-AEP, while the model diatom *Phaeodactylum tricornutum* failed [17]. However, our recent study showed that P-starvation treated *P. tricornutum* can recover the growth with 2-AEP supplement, and the meta-omic atlas suggest that the associated celluar mechanism is prevalent in diatom assemblages and more active in the cold regions [16]. Further exploration is required to address the discrepancies and ecological implication with regards to the biogenic phosphonate utilization by diatom.

Diatoms are important eukaryotic phytoplankton with diverse functional resilience that contribute ∼20% of global carbon fixation and 40% of marine primary productivity [18]. Diatom distribution and abundance are of great importance to marine biogeochemical cycling due to the ballast effect attributed to the efficient settling of siliceous cell shells [19]. Plankton genomics-based model prediction shows that sea surface temperature and phosphate are two main drivers of the plankton community under climate change, and projected decrease of carbon export fluxes are associated with diatoms [20]. Therefore, sorting out the variable stoichiometry of diatoms under different phosphorus conditions and the effects of temperature will provide further insight of the role of diatoms in the biogeochemical cycle.

In our previous study, we reported that model diatom *Phaeodactylum tricornutum* is able to utilize 2-AEP as an alternative P source, and the metabolism-associated gene homologs distribute globally in temperature-dependence, with a high proportion derived from *Thalasiosira pseduonana* [16]. *T. pseudonana* is also a model diatom species widely studied for its dominancy in natural phytoplankton assemblage [21-23].

Promoted by that, here we examine the bioavailability of both 2-AEP and 2-AP3 by the cosmopolitan diatom species *T. pseudonana*. We further investigate the cell physiological response and cellular stoichiometric ratio of the cultures grown under different P conditions and temperatures as well. All these findings fill the puzzle piece to the overall picture of the marine P biogeochemical cycle.

## Materials and methods

### Cell culture and experiment setup

*T. pseudonana* used in this study was provided by Center for Collections of Marine Bacteria and Phytoplankton of Xiamen University. 2-AEP and 2-AP3 were obtained from Sigma-Aldrich (St. Louis, MO, USA). The algal strain was cultured in filtered natural seawater enriched with f/2 medium. Two batch cultures were grown at different temperatures and phosphorus conditions, and antibiotics were applied to inhibit the growth of bacteria, each group in triplicate (Table 1). Before the experiment, seed cultures of *T. pseudonana* were subjected to phosphorus starvation for 8∼10 days until the phosphate concentration in the medium was below ∼0.3 μM and the cell growth ceased. The aim of Batch 1 culture is to examine the bioavailability of 2-AEP and 2AP3, and the aim of Batch 2 culture is to further explore the cellular physiological response to 2-AEP.

**Table 1.** Culture conditions (a,b: control groups in Batch 1, b: control group in Batch 2)

| Culture | P nutrient concentration | T (°C) | Culture condition |
| --- | --- | --- | --- |
| **Seed** | Starvation treated<br>8-10 day (<0.3 µM) | 20 | • f/2 medium (salinity=30)<br>• 14:10 light: dark cycle |
| **-P<sup>a</sup>** | No addition |  | • photon flux: 180 µmol m<sup>-2</sup> s<sup>-1</sup> |
| **+P<sup>b</sup>** | DIP (36 µM) |  | • Antibiotics cocktail (final concentration in medium: ampicillin 100 mg/L, streptomycin 50 mg/L and kanamycin 50 mg/L) |
| **Batch 1** | 2-AEP (36 µM)<br>2-AP3 (36, 72 µM) |  |  |
| **Batch 2** | 2-AEP (72 µM) | 12/25 |  |

### Determination of cell density and Fv/Fm

The cell density was measured daily by using a CytoFLEX flow cytometer (Beckman Coulter, Indianapolis, IN, USA), as estimated by gating areas in the Chlorophyll A vs SSC-A dot plot generated from a 1 mL cell sample. Fv/Fm was determined by using a FIRe Fluorometer System (Satlantic, Halifax, NS, Canada). Prior to the measurement, 1 mL cell sample was subjected to dark adaption for 20 mins and processed following the manufacture protocol [24].

### Cellular carbon, nitrogen and phosphorus content measurement

Collected cell samples were pretreated as described before [16]. Cellular carbon and nitrogen contents were determined using a vario EL cube analyzer (Elementar Analysensysteme GmbH, Hanau, Germany) followed reported method [25]. Regarding cellular phosphorus determination, cells filtered onto GF/F membrane were digested by autoclaving at 121 °C, 30 min, then total phosphorus was determined by the molybdenum method [26]. The ratio of C:P and N:P was obtained by calculation.

## Results and discussions

Our previous study proposed that a putative 2-AEP utilization pathway involved with endocytosis and membrane incorporation is ubiquitous in diatom assemblages [16]. 2-AP3 is the N-methyl derivative deviate of 2-AEP and a membrane constituent (Fig. 1) [11]. Therefore, we propose the hypothesis that dominant diatom species can utilize these two biogenic phosphonates as well. As expected, our results confirm this hypothesis as elaborated below.

**Fig. 1.**
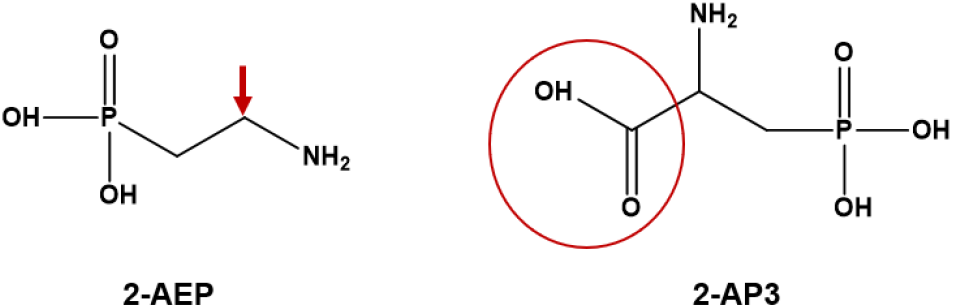
Chemical structural of 2-AEP and 2-AP3. Arrow denotes the α-position of 2-AEP, and the circle denotes the -COOH added in the α-position of 2-AP3.

### Promoted cell growth by 2-AEP and 2-AP3 showing different patterns

In batch 1 culture, we observe different growth-promoting effects between the 2-AEP and 2-AP3 groups. After phosphorus starvation, *T. pseudonana* was able to recover growth significantly in the medium supplied with 2-AEP (36 μM) and 2-AP3 (72 μM) respectively (P<0.05), while failing in 2-AP3 supplement (36 μM) (Fig. 2a).

**Fig. 2.**
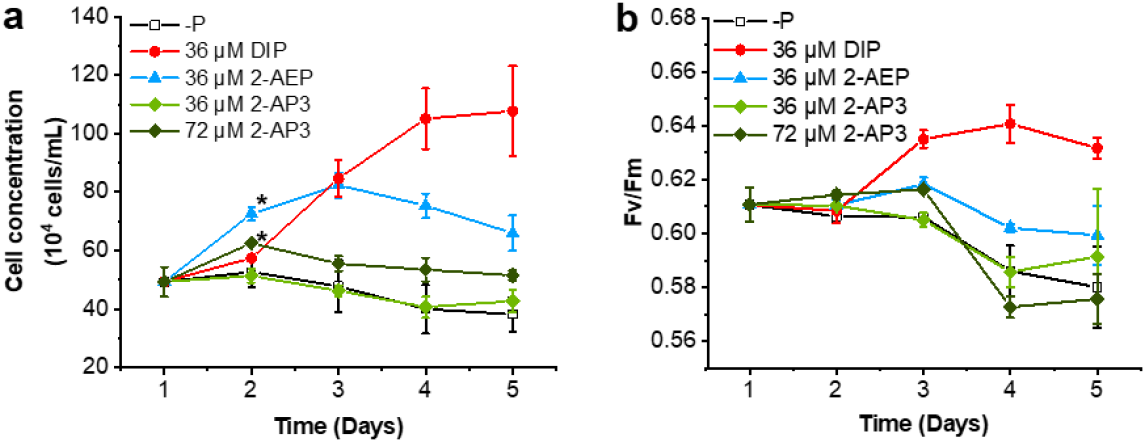
Physical response of *T. pseudonana* to different phosphorus. (a) Growth curve. (b) Fv/Fm. Each group were triplicate, * P<0.05.

Cells in the 2-AEP (36 μM) group continued growth and peaked the maximum cell concentration of 8.24×10^5^ cells mL^-1^ at Day 3, then declined gradually. In consistent with that, Fv/Fm increased slightly in the first three days, then declined along with the cell density (Fig. 2b).

In comparison, we observed mild growth promotion in the 2-AP3 group. After supplied with 72 μM 2-AP3, instant cell growth was recovered in 24 h, showing growth rate 0.24 μ d^-1^ about half of that in the 2-AEP (36 μM) group (Fig. 2a). After then, cell ceased growth and declined slightly till the end of this experiment, accompanied by abrupt decline of Fv/Fm (Fig. 2b). Maximum cell density observed on D2 was 6.26×10^5^ cells mL^-1^, about half of that in the 2-AEP (36 μM) group and 2-fold of that in the DIP-replete group.

These results demonstrate that both 2-AEP and 2-AP3 can be utilized as an alternative phosphorus source to support the cell growth of *T. pseudonana*. Though provided in the same or higher concentration, lower cell density acquired in both phosphonate groups suggest limited utilization efficiency of these compounds compared with the P-replete group. Lower biomass acquired in phosphonate cultures is common when compared to equivalent DIP culture [15, 17]. Furthermore, lower Fv/Fm indicates repressed photosynthesis, suggesting *T. pseudonana* cells were under phosphorus stress in both phosphonate groups.

In our previous study, we propose that 2-AEP is transported into diatom cells through endocytosis and then incorporated into the phospholipid of cell membrane [16]. 2-AP3 is the N-methyl derivatives deviate of 2-AEP, known as the component of cell membrane as well. Studies have demonstrated that 2-AEP and its N-methyl derivatives can be incorporated into phospholipids in cell membrane [27-29]. *P. tricornutum* is fail to recover growth after supplied with 2-AP3 (Fig. S1). Despite that, the previous mechanism deduced from *P. tricornutum* might apply to explain the promotion of *T. pseudonana* cell growth by 2-AP3 in this study. It is found that 2-AP3 can be decomposed via transamination reaction and decarboxylation to 2-AEP in *Tetrahymena* [30]. Given the limited utilization of 2-AP3 observed in this study, we cannot completely exclude the other possible metabolic pathway. The underlying mechanism of 2-AP3 utilization by *T. pseudonana* and its bioavailability by other phytoplankton need to be further investigated to address the knowledge gap in eukaryotic phytoplankton.

### Different growth strategies in the 2-AEP and DIP cultures under low temperature

Through data mining in the global ocean atlas, we found enriched distribution of representative genes of the proposed mechanism in lower temperature waters [16]. Therefore, we conducted the second batch culture to further explore the effect on cell growth while supplied with 2-AEP under different temperatures. In this experiment, 2-AEP was provided in a higher concentration 72 μM to achieve optimal growth results based on pre-experiment.

Out of our expectation, we observed entirely different growth pattern in both DIP-replete and 2-AEP (72 μM) under different temperatures (Fig. 3). Generally, *T. pseudonana* cells exhibited a sustained growth at 12 °C with higher cell density, but a short-time rapid growth at 25 °C with lower cell density, regardless of phosphorus condition. At 12 °C, the cell density barely changed in the first 24 h, then increased steadily in both 2-AEP and DIP groups till D6, sharing no difference in the first 4 days (Fig. 3a). While cultured at 25 °C, cell density exhibited rapid growth in the 48 h then entered the stationary phase after D4 in the DIP group, and cells ceased growth after D3 in 2-AEP group then declined significantly. Comparably, it has indicated that the growth rate of *T. pseudonana* is higher under 8∼17 °C than that under 17∼25 °C with sufficient DIP supply [31]. We also observe such a manner in our study of *P. tricornutum*, cells grow continuously at 12 °C and a short-term rapid growth at 25 °C (data unpublish). Such a difference can be explained by the classical *r*-/*K*-selection paradigm, in which *r*-strategist is featured with high growth rate as opportunistic and *K*-strategist is described as equilibrium [32].

**Fig. 3.**
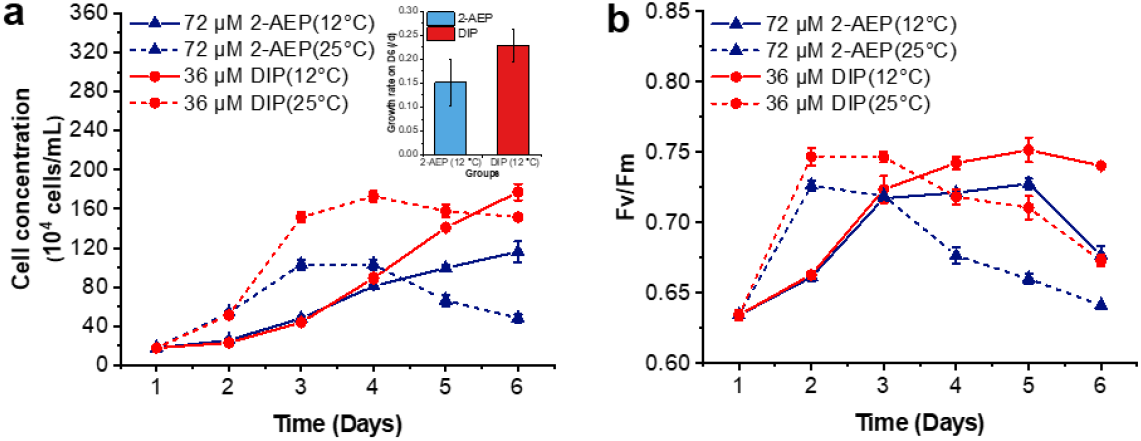
The effect of temperature on *T. pseudonana* physiological parameters under different phosphorus conditions, (a) cell concentration, inner panel represents the growth rate on D6 at 12 °C, (b) Fv/Fm.

Besides that, we notice that there is a significant difference in the promotion effect under different culture conditions (Fig. 3a). In the DIP-replete groups, the final cell concentration (177.1 ± 8.4 × 10^4^ cells mL^-1^) grown at 12 °C is slightly higher than that (151.7 ± 2.7 × 10^4^ cells/mL) in the 25 °C culture. But in 2-AEP groups, the final cell concentration (116.2 ± 10.9 × 10^4^ cells/mL) in the 12 °C culture is notably higher than that (48.2 ± 3.9 × 10^4^ cells/mL) in the 25 °C culture, about 3 times as much. When cultured at 12 °C, the growth rate on D6 of 2-AEP group is 0.152 ± 0.049 μ d^-1^, about 66.4% of that in the DIP-replete group (Fig. 3a).

Variations of Fv/Fm showed a comparable pattern in accordance with the growth curve (Fig. 3b). Dramatic increase of Fv/Fm values in the first 24 h indicate that instant recovery of photosynthetic capacity accounts for the rapid growth in both cultures under 25 °C. After day 3, Fv/Fm values declined continuously to the initial value, consistent with the cells entering stationary phase in 25 °C cultures. In contrast, to meet the sustained cell growth under 12 °C, Fv/Fm value increased steadily, higher in the DIP-replete group than that in 2-AEP group.

Taken our results together, we conclude that *T. pseudonana* cells cultured with 2-AEP exhibit better physiological adaptability as *K*-strategist under lower temperature, result in a significant elevated cell density, even comparable to that in the DIP-replete cultures. Meanwhile, Fv/Fm value represents a good consensus of higher photosynthetic capacity and a higher growth rate.

### Changes of cellular elemental stoichiometry of *T. pseudonana*

Over decades, Redfield ratio 106C:16N:1P (Redfield 1958) has been widely accepted as an index of phytoplankton cellular elemental stoichiometry, and used to assess the nutrient status of phytoplankton [33]. For instance, offset environmental C:P and N:P ratios (C:P higher than 106:1 and N:P higher than 16:1) is assumed as phosphorus limitation for phytoplankton growth [34, 35]. However, many studies have revealed that marine phytoplankton elemental stoichiometric ratios deviate from that empirical ratio, thus playing a major role in shaping the environmental stoichiometry ratio [36, 37].

#### Stoichiometry variation under different phosphorus conditions

In this study, the N:P ratio of DIP-replete group (∼6:1) (Fig.4a) is far below that of Redfield ratio and group-specific optimal value 14:1 [38]. This can be explained by the luxury uptake of DIP and storage in the form of polyP after phosphorus starvation, which is typical in diatoms [39, 40]. Regarding the initial phosphorus starvation state, the C:P and N:P ratios were 117.72 ± 23.14 and 24.25 ± 7.94 respectively, in consistent with reported cellular stoichiometry in diatom under insufficient nutrient condition [41]. After that, the C:P and N:P ratios decreased significantly to 36.93 ± 3.96 and 5.63 ± 0.56 respectively on D5 in the DIP-replete group, while continue increase or maintain under other different phosphorus conditions (Supplementary Fig. S2a).

**Fig. 4.**
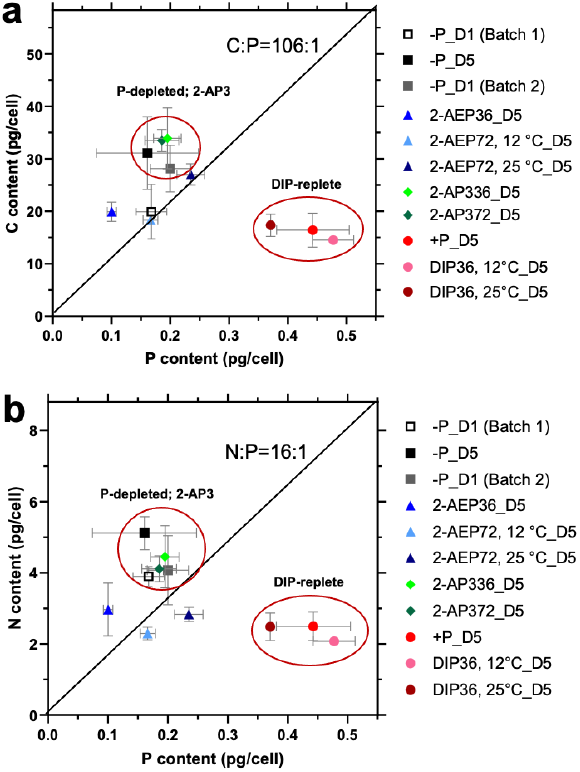
Cellular stoichiometry of *T. pseudonana* under different conditions. (a) C:P ratio; (b) N:P ratio.

Stoichiometry of 2-AP3 groups (36 μM and 72 μM) is similar to that of the P-depleted group (Fig. 4), in consistent with the growth pattern. P-depleted cells and cells treated with 2-AP3 were grouped together showing higher C:P and N:P ratios, mainly because of higher cellular carbon and nitrogen content, as well as lower cell phosphorus content than that in the DIP-replete groups (Fig. S3a).

#### Effect of temperature on cellular stoichiometry

In the DIP-replete groups, there is no significant difference identified between 12 °C and 25 °C (Fig. 4, Supplementary Fig. S2b). In 2-AEP group, the C:P and N:P ratios were lower than that in P-depleted *T. pseudonana* and higher than that in DIP-replete group. In 72 μM 2-AEP cultures (12 °C and 25 °C), the C:P and N:P values were 111.2 ± 11.1 ∼ 115.8 ± 12.1 and 13.9 ± 1.8 ∼ 12.1 ± 0.6 respectively, closer to the Redfield ratio with higher 2-AEP concentration and lower temperature.

The interactions between environment conditions and cell growth are the key factors to drive stoichiometric variation [37], temperature is the major factor due to the direct effect on cell growth [36, 42]. Global research on phytoplankton stoichiometry has found that both C:P and N:P ratios decrease with temperature [36, 43, 44], but laboratory evidence is insufficient regarding species difference. Our findings firstly show the synergetic effect of temperature and phosphonate on cellular stoichiometry in *T. pseudonana*, higher N:P ratios when cultured with 2-AEP under lower temperature. In the DIP-replete group, the lower temperature significantly decreased the C:P and N:P ratios of *T. pseudonana*, in consistent with previous report [36, 45, 46].

It is reported that cells increase ribosomes concentration and cellular P content to compensate for low translation efficiency of ribosomes at low temperature [47]. Such a hypothesis is confirmed with observation of strong temperature dependency of C:P and N:P in high latitude ecosystems [48]. In this study, the cellular P content of DIP-replete groups was significantly higher at 12 °C compared with 25 °C (Fig. S3b), indicating that low temperature might increase the phosphorus demand and promote the phosphorus absorption of *T. pseudonana*. In addition, temperature also had significant effect on the C:P and N:P ratios in DIP group. In the 2-AEP group, barely temperature effects were observed on stoichiometry of *T. pseudonana*. This might attribute to the stable chemical properties of 2-AEP, hard to be hydrolyzed into phosphate.

### High variability of stoichiometry in diatom

Nutrient availability is considered to be the major driver shaping the phytoplankton stoichiometry, for example, phosphorus limitation accounts for the increase of C:P and N:P ratios of phytoplankton [49, 50]. In this research, the C:P and N:P ratios of *T. pseudonana* declined rapidly after 36 μM DIP supplement. Both C:P and N:P ratios are higher than the Redfield ratio under DIP-depleted condition, whereas significantly lower than the Redfield ratio under P-replete condition. Compared with other studies, the C:P and N:P in this study was much lower no matter under P abundant or P deficiency conditions (Fig. 5) [36, 50-52]. In agreement with the previous results, the C:P and N:P ratios of DIP-depleted *T. pseudonana* were higher than the Redfield ratio [50, 53, 54]. According to the observation data from ALOHA and BATS stations, C:P ratio of suspended particles varies mostly from 100 to 200 while below the P-stressed threshold [55]. Our results show that *T. pseudonana* grown with 2-AEP are close to the classical Redfield ratio.

**Fig. 5.**
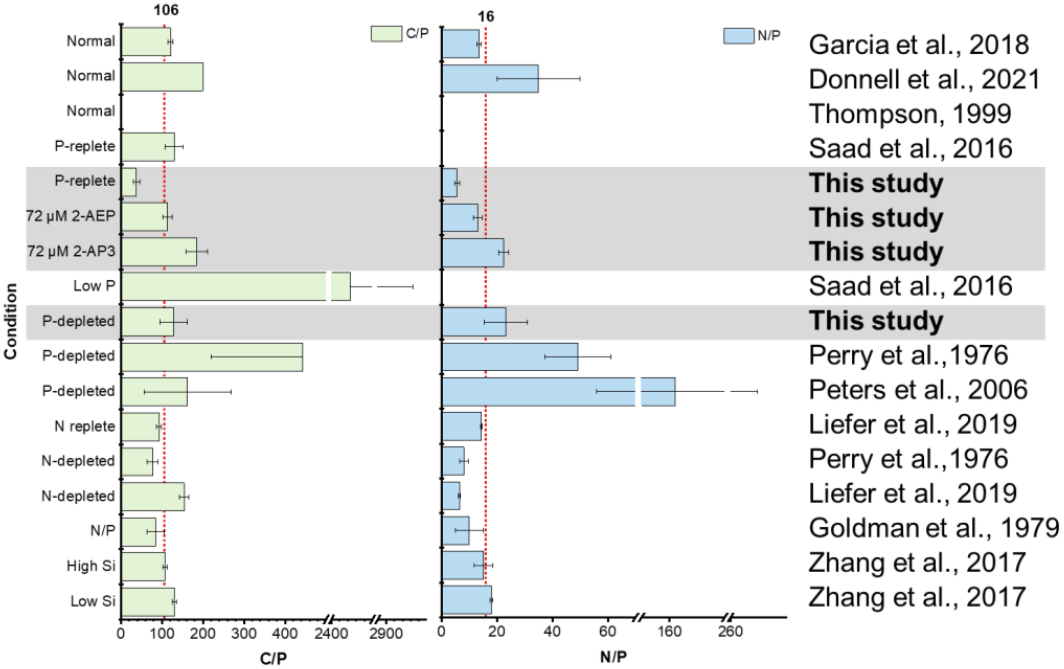
Comparison of elemental stoichiometry of *T. pseudonana* in this study and previous research.

The effects of other nutrients such as nitrogen or silicon have been studied as well. N deficiency lead to lower N:P [53, 56, 57] and silicon concentration has no significant effect on both C:P and N:P (Fig. 5) [58]. Though summarizing and comparing with previous reports, our study provides useful information of temperature and P nutrient effects on stoichiometry variation in *T. pseudonana*.

## Summary and Ecological implications

Overall, our study has three major findings. (1) It demonstrates that biogenic phosphonate 2-AEP and 2-AP3 can be utilized by diatom *T. pseudonana* to support cell growth, and 2-AEP is more preferable than 2-AP3 evidenced by higher cell density. (2) Disparate growth strategies are identified under different temperatures, and significantly promoted cell growth of 2-AEP culture under lower temperature compared to mild temperature indicates its adaptive function. (3) The mediate value of C:P and N:P ratios in 2-AEP groups between that in P-depleted (2-AP3) and DIP-replete groups suggest the potential effect on environmental elemental stoichiometry may follow.

Such a scenario has importing ecological implications in sub-polar region, diatoms represent the major primary producer. In the regions of phosphorus deficiency, species like *T. pseudonana* has a significantly advantage over other species which unable to use phosphonates. Our result is consistent with reported optimal growth rate of diatom is 15?C [59]. As stated in the predicted future paradigm, P-stress conditions in the surface water will be intensified due to the reduced nutrient supply from bottom water as a result of global warming [55]. Statistical model also suggests that C:P and N:P ratios will increase at high latitudes in the future [48]. Phytoplankton with P nutrient uptake plasticity are critical for the ocean primary production [55]. Element stoichiometry might be altered along with the variation of phytoplankton community composition. Different species contributing disparate to the flux of particulate organic carbon (POC) [60]. Our findings of 2-AEP and 2-AP3 bioavailability by *T. pseudonana* and the stoichiometry variation under different phosphorus and temperature conditions provide new insights in understanding the diatom ecophysiology in marine biogeochemical cycling and the effect on marine organic matter stoichiometry as well.

## Acknowledge

This research was supported by National Key R&D Program of China grant 2022YFC3105302 (X.L.), and Natural Science Foundation of Fujian Province of China, # 2020J06008, for Distinguished Young Scholars (J.M.). We thank Junhui Chen for the assistant of cellular carbon and nitrogen determination.

## Notes

### Competing Interest Statement

The authors have declared no competing interest.

